# The *C. elegans* uv1 neuroendocrine cells provide direct mechanosensory feedback of vulval opening

**DOI:** 10.1101/2024.04.11.589042

**Authors:** Lijie Yan, Alexander Claman, Addys Bode, Kevin M. Collins

## Abstract

Neuroendocrine cells react to physical, chemical, and synaptic signals originating from tissues and the nervous system, releasing hormones that regulate various body functions beyond the synapse. Neuroendocrine cells are often embedded in complex tissues making direct tests of their activation mechanisms and signaling effects difficult to study. In the nematode worm *C. elegans*, four uterine-vulval (uv1) neuroendocrine cells sit above the vulval canal next to the egg-laying circuit, releasing tyramine and neuropeptides that feedback to inhibit egg laying. We have previously shown uv1 cells are mechanically deformed during egg laying, driving uv1 Ca^2+^ transients. However, whether egg-laying circuit activity, vulval opening, and/or egg release triggered uv1 Ca^2+^ activity was unclear. Here we show uv1 responds directly to mechanical activation. Optogenetic vulval muscle stimulation triggers uv1 Ca^2+^ activity following muscle contraction even in sterile animals. Direct mechanical prodding with a glass probe placed against the worm cuticle triggers robust uv1 Ca^2+^ activity similar to that seen during egg laying. Direct mechanical activation of uv1 cells does not require other cells in the egg-laying circuit, synaptic or peptidergic neurotransmission, or TRPV and Piezo channels. EGL-19 L-type Ca^2+^ channels, but not P/Q/N-type or Ryanodine Receptor Ca^2+^ channels, promote uv1 Ca^2+^ activity following mechanical activation. L-type channels also facilitate the coordinated activation of uv1 cells across the vulva, suggesting mechanical stimulation of one uv1 cells cross-activates the other. Our findings show how neuroendocrine cells like uv1 report on the mechanics of tissue deformation and muscle contraction, facilitating feedback to local circuits to coordinate behavior.

**Significance Statement:** Neuroendocrine cells respond to diverse physical and chemical signals from the body, releasing hormones that control reproduction, gut motility, and fight or flight responses. Neuroendocrine cells are often found embedded in complex tissues, complicating studies of how they are activated. Using the genetic and experimental accessibility of *C. elegans*, we find the uv1 neuroendocrine cells of the egg-laying motor behavior circuit respond directly to mechanical stimulation and vulval opening. We show that L-type voltage-gated Ca^2+^ channels facilitate uv1 mechanical activation and also coordinate cell activation across the vulval opening. In contrast to other mechanically activated cells, uv1 activation does not require Piezo or TRPV channels. This work shows how neuroendocrine cells relay critical mechanosensory feedback to circuits that control reproduction.

## Introduction

How activity in neural circuits is coordinated to drive animal behaviors is a fundamental question in neuroscience. Neuroendocrine cells play essential, but understudied, roles in neural circuits. Neuroendocrine respond to physical, chemical, and synaptic signals from both tissues and the nervous system, produce action potentials and secrete hormones that act extrasynaptically to control body functions (Watson and Breedlove, 2016). For example, Enterochromaffin Cells in the gastrointestinal system respond to physical stretch via mechanoreceptors including Piezo channels, releasing serotonin that has a myriad of physiological functions (Alcaino et al., 2018; Linan-Rico et al., 2016). Pulmonary neuroendocrine cells in the lung respond to fluctuations in oxygen levels and release bioactive neuropeptides (Bordon, 2016). Disruption of neuroendocrine cell function can cause human disease. For example, dysfunction of pulmonary neuroendocrine cells is related to several debilitating airway diseases including acute hypoxia, asthma, and congenital diaphragmatic hernia (Bordon, 2016; Cutz et al., 2007). How neuroendocrine cells respond to sensory and mechanosensory signals, and how this regulates secretion of neurotransmitters and neuropeptides that modulate downstream circuits and physiology, is not well understood. Neuroendocrine cells express mechanosensitive ion channels that contribute to their mechanical activation (Alcaino et al., 2017), but direct genetic tests exploring which physiologically relevant signals contribute to neuroendocrine cell activation and signaling are lacking.

Since many neuroendocrine cells are found embedded in complex tissues, direct experiments to manipulate and investigate the function of these cells can be technically challenging. The uv1 neuroendocrine cells of the *C. elegans* vulval egg-laying apparatus provide a convenient, genetically accessible model to investigate conserved features of neuroendocrine cell signaling. Egg-laying behavior of *C. elegans* is driven by a simple neural circuit (Collins et al., 2016). A pair of Hermaphrodite-Specific command Neurons (HSNs) synapse onto vulval muscles, releasing serotonin and neuropeptides that stimulates the contractility of the postsynaptic vulval muscles (Brewer et al., 2019; Olson et al., 2023). The coordinated contraction of vulval muscles vm1 and vm2 open the vulva and allow egg release (Brewer et al., 2019; Li et al., 2013). The vulval muscles are also innervated by the female-specific cholinergic Ventral C (VC) motor neurons (Pereira et al., 2015) which are mechanically activated during by muscle contraction, facilitating serotonin-induced egg laying and its coordination with locomotion (Collins et al., 2016; Kopchock et al., 2021).The circuit also has a set of neuroendocrine cells, the uv1s, that are mechanically deformed and subsequently activated following passage of eggs through the vulva and egg release (Collins et al., 2016). Once activated, the uv1 cells release tyramine and neuropeptides that signal to inhibit HSN activity and terminate the egg-laying behavior active state (Alkema et al., 2005; Banerjee et al., 2017; Collins et al., 2016). Whether uv1 activation is direct, or instead mediated by other, rhythmic and sequential activity in the egg-laying circuit (Medrano and Collins, 2023), has not been explored. uv1 expresses three different TRPV channels that might function as heteromers to facilitate their mechanical activation (Jose et al., 2007; Upadhyay et al., 2016), but there have been no direct tests of their function in uv1 mechanical activation.

Here, we developed optogenetic and direct prodding methods to investigate how the uv1 cells are mechanically activated. We found that optogenetic stimulation of vulval muscle contraction, bypassing ongoing egg-laying circuit activity, triggers uv1 mechanical activation independent of egg passage. Direct mechanical prodding of uv1 similarly induced a strong and reproducible Ca^2+^ transient in uv1 cells. We show this mechanical activation of uv1 cells is independent of other cells in egg-laying circuit, neurotransmission, and candidate mechanically activated channels including TRPV and Piezo (Bai et al., 2020; Jose et al., 2007). We find the L-type Ca^2+^ channel EGL-19 promotes uv1 activation independent of other voltage-gated and store-operated channels. L-type Ca^2+^ channels also facilitate the coordinated activation of uv1 cells across the vulval opening, suggesting uv1 mechanical interactions *in trans* communicate feedback to the rest of the egg-laying behavior circuit.

## Results

### uv1 cells are mechanically activated in response to egg release

Egg-laying behavior of *C. elegans* is driven by a simple neural circuit (Fig. 1A) (Collins et al., 2016). The uv1 cells sit at the dorsa-lateral surface of the vulval canal, adjacent to the HSN and VC presynaptic termini that release neurotransmitters onto the vm2 vulval muscle arms (Collins and Koelle, 2013; Ravi et al., 2018a). We developed ratiometric Ca^2+^ imaging methods to record *in vivo* cell activity in the circuit by using the calcium reporter GCaMP5 co-expressed with the Ca^2+-^insensitive red fluorescent protein mCherry (Ravi et al., 2018b). Our previous uv1 Ca^2+^ reporter based on the *ocr-2* promoter/enhancer expressed rather weakly in the uv1 cells and with strong additional co-expression in head and tail sensory neurons along with the uterine seam (utse) cells (Collins et al., 2016; Jose et al., 2007). Combining the *tdc-1* promoter (Alkema et al., 2005) with the *ocr-2* 3’ untranslated region (see methods), we improved the signal to noise of our uv1 Ca^2+^ imaging while reducing reporter expression in other cells. Ratiometric Ca^2+^ imaging with this new reporter in behaving worms shows the uv1 neuroendocrine cells were mechanically deformed by egg release leading to strong Ca^2+^ transient activity (Fig. 1B-C), as previously shown with the previous reporter (Collins et al., 2016).

**Figure 1:**
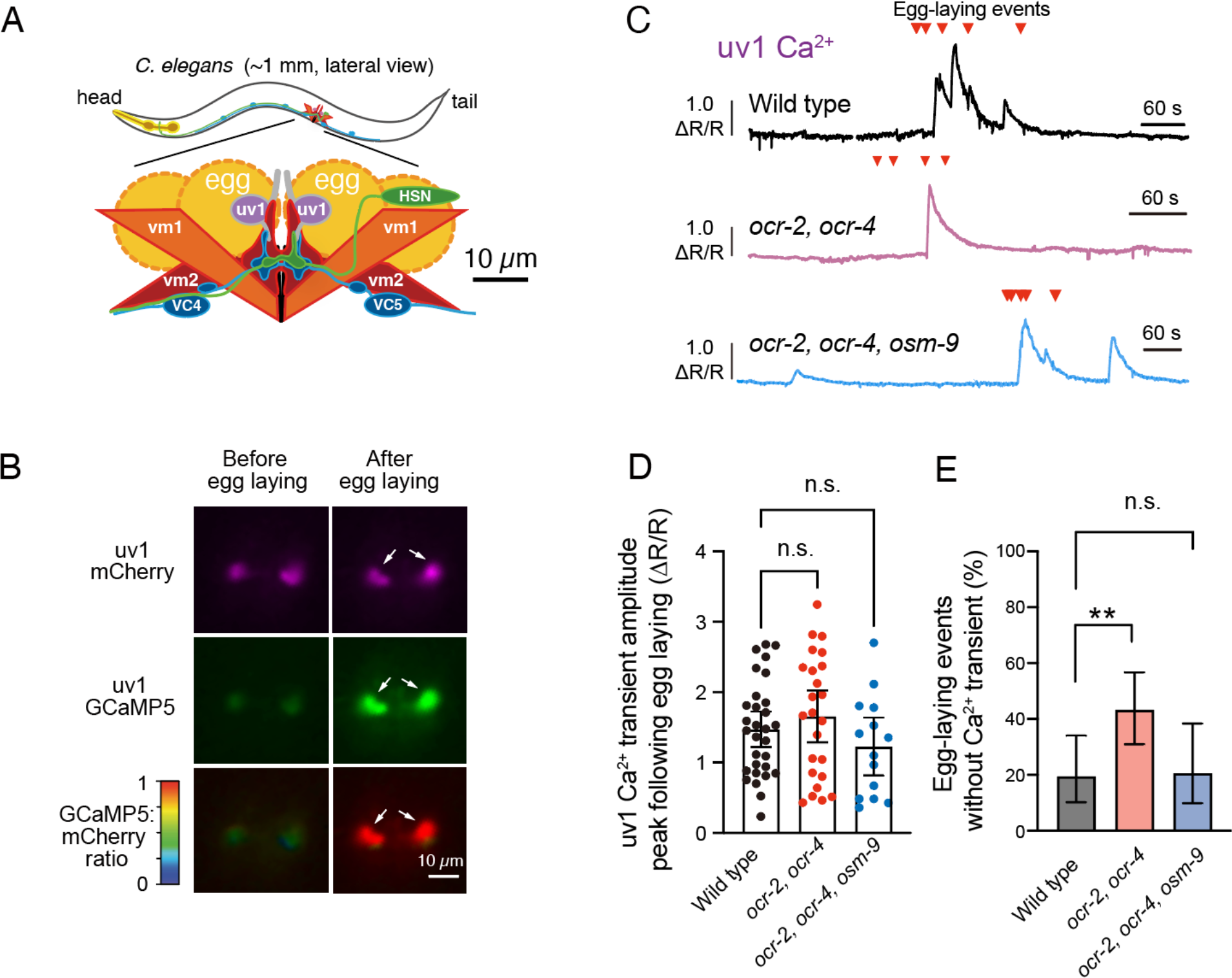
uv1 cells are mechanically activated in response to egg release. **(A)** Graphical representation of egg-laying circuit in *C. elegans* (Collins et al., 2016). **(B)** Micrographs of uv1 GCaMP5 and mCherry fluorescence and the GCaMP5/mCherry ratio before and after egg laying. Arrowheads indicate uv1 position and increase in GCaMP5 fluorescence; scale bar, 10μm. **(C)** Representative traces (10 min) of the uv1 GCaMP5/mCherry ratio in wild-type (black), *ocr-2 ocr-4* (red), and *ocr-2 ocr-4 osm-9* (blue) TRPV channel mutant animals; egg-laying events are indicated by arrowheads. **(D)** Scatterplots of uv1 Ca^2+^ transient amplitudes following egg-laying event in wild type (black), *ocr-2, ocr-4* (red), and *ocr-2, ocr-4, osm-9* (blue) mutant animals. Points represent each transient from 10 animals; bar indicates mean from X animals; n.s. represents p >0.05; one-way ANOVA with Bonferroni’s correction for multiple comparisons. **(E)** Bar graphs (±95% confidence interval for the proportion) comparing percentage of egg-laying events that fail to induce uv1 Ca^2+^ activity in wild-type (black), *ocr-2, ocr-4* (red), *ocr-2, ocr-4, osm-9* (blue). ** indicates p <0.01 (Fischer’s exact test).

Mechanosensitive TRPV channel subunits OCR-2, OCR-4, OSM-9 have previously been shown to be expressed in the uv1 cells (Jose et al., 2007). These heteromeric channels are thought to promote cell excitability (Upadhyay et al., 2016) and tyramine and release (Jose et al., 2007) to inhibit egg-laying behavior (Collins et al., 2016). To better understand how TRPV channels regulate uv1 Ca^2+^ activity during egg release, we performed ratiometric Ca^2+^ imaging in freely behaving *ocr-2(ak47), ocr-4(vs137)* double mutants and *ocr-2(ak47), ocr-4(vs137), osm-9(cx10)* triple mutants alongside wild-type control animals (Fig. 1C). We found that egg-laying events still triggered strong uv1 Ca^2+^ activity in TRPV double and triple mutants (Fig. 1D-E). Quantitation of uv1 Ca^2+^ transient peak amplitudes following egg-events were not significantly different (Fig. 1F), suggesting no gross defects in uv1 cell activation. As seen in wild-type animals, not all egg-laying events in TRPV mutants induced a Ca^2+^ transient. While there was a significant increase in this failure rate in *ocr-2, ocr-4* double mutants, this was not shared in the *ocr-2, ocr-4, osm-9* triple mutant animals (Fig. 1G). These data show that while TRPV channels may regulate uv1 responses to egg-laying events, they are not strictly required.

### Optogenetic vulval muscle stimulation triggers uv1 Ca^2+^ activity in response to vulval opening but not egg release

Egg laying in *C. elegans* is a two-state behavior where 3-5 eggs are laid within ∼2 min active states separated by ∼20 min inactive states where no eggs are laid (Waggoner et al., 1998). As uv1 Ca^2+^ transient activity occurs when other cells in the egg-laying circuit are active (Collins et al., 2016), whether vulval opening and/or egg release mechanically activates uv1 directly or if the uv1 cells instead respond to signaling from other cell(s) in the circuit (or both) is not clear. To address whether vulval opening and/or egg release are sufficient to activate the uv1 cells, we transgenically expressed Channelrhodopsin-2 (ChR2) in the vulval muscles which allows for the optogenetic stimulation of the vulval muscles and egg release following exposure to blue light (Kopchock et al., 2021). As above, we co-expressed GCaMP5 and mCherry in the uv1 cells to record changes in uv1 Ca^2+^ activity following optogenetic vulval muscle stimulation (Fig. 2A). Vulval muscle stimulation was sufficient to trigger a rapid separation of the uv1 cells (Fig. 2B) which was accompanied by a sharp increase in uv1 cell Ca^2+^ activity (Fig. 2D) and egg laying. Analysis of distance between anterior and posterior uv1 cells showed a clear and significant separation starting ∼2 s before egg release, confirming that optogenetic stimulation of the vulval muscles drove contraction (Fig. 2C). Interestingly, uv1 separation decreased almost immediately upon egg release, despite ongoing vulval muscle optogenetic stimulation (Fig. 2D). To test whether egg release was required for uv1 activation, we sterilized animals with Floxuridine (FUDR), which prevents generation of embryos (Mitchell et al., 1979). Importantly, sterilization does not impede optogenetic activation of the vulval muscles (data not shown). Optogenetic vulval muscle stimulation induced robust uv1 Ca^2+^ activity, even in sterilized animals (Fig. 2D). These results show that uv1 mechanically responds to vulval opening, not by the deformation or passage of eggs *per se*, and is separable in part from circuit activity normally associated with the egg-laying active state.

**Figure 2:**
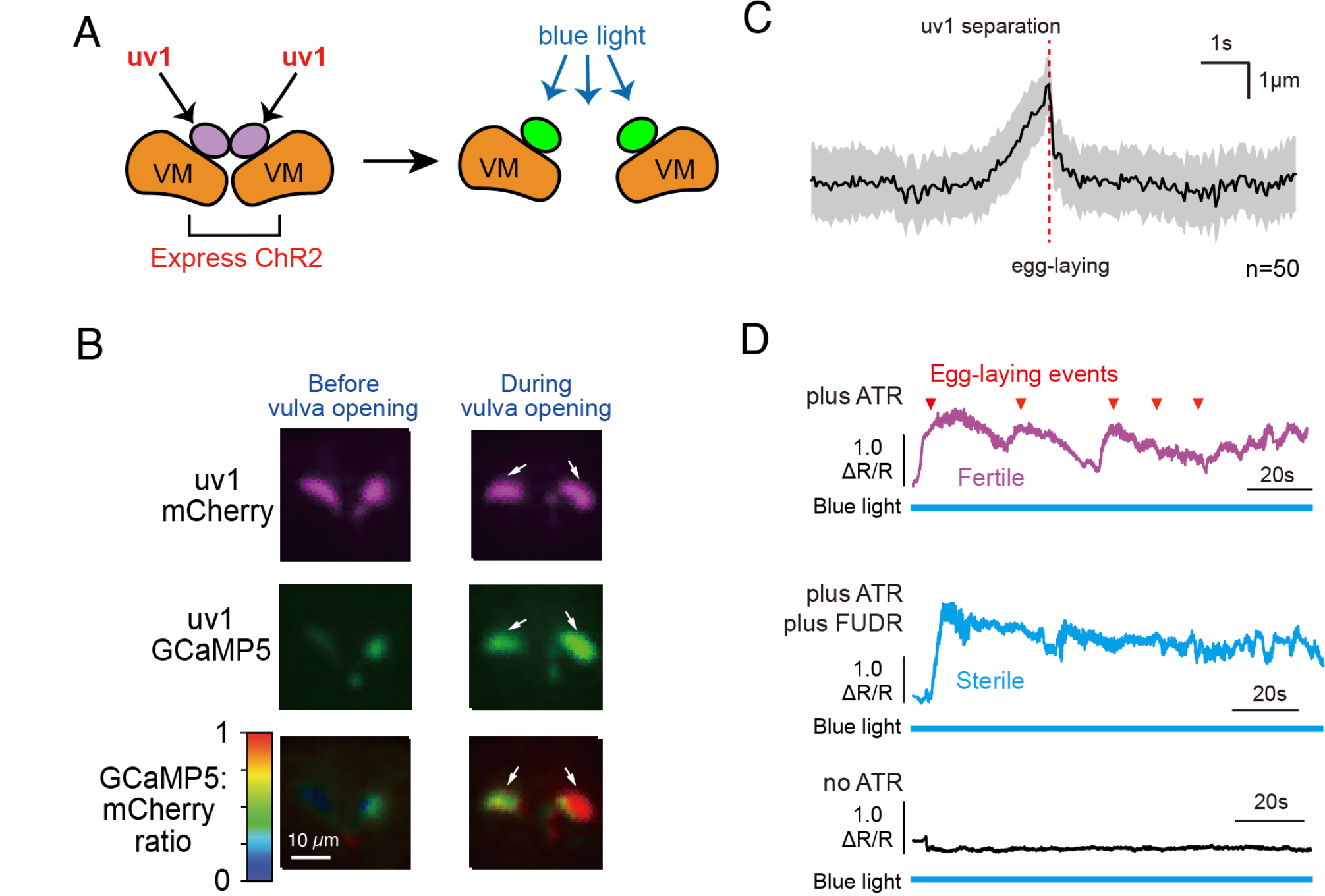
Optogenetic stimulation of the vulval muscles drives uv1 activation in response to vulval opening. **(A)** Diagram of optogenetic method with Channelrhodopsin-2 expressed in vulval muscles (orange) and GCaMP5/mCherry in the uv1 cells (purple). Arrowheads represents blue light exposure and predicted uv1 GCaMP5 Ca^2+^ response (green). **(B)** Micrographs of uv1 Ca^2+^ transient activity before and during ChR2 activation of vulval muscles and egg laying. Arrowheads indicates position of uv1 cells following contraction and vulval opening. **(C)** Average distance between anterior/posterior uv1 cells 5 s before and after egg-laying events (n=50 egg-laying events from 13 animals). Grey area indicated 95% confidence interval for the mean. Red dotted line indicates when eggs got laid. **(D)** GCaMP5/mCherry ratio (ΔR/R) trace of uv1 Ca^2+^ activity following ChR2 activation in animals with all-*trans* retinol (ATR; top, purple), with both ATR and following FUDR sterilization (middle, blue), and in animals grown without ATR, a necessary cofactor for ChR2 activity or FUDR (bottom, black). Egg-laying events are indicated by red arrowheads. Blue horizontal lines represent continuous blue light exposure.

### The uv1 cells are activated by direct mechanical stimulation

While optogenetic stimulation of the vulval muscles bypasses activity in some cells in the egg-laying circuit (Dhakal et al., 2022), it also induces activity in others like the VC neurons (Kopchock et al., 2021). Vulval muscle activity can also promote downstream Ca^2+^ transient activity in other cells of the circuit including the HSNs (Medrano and Collins, 2023; Ravi et al., 2018a). To test whether direct stimulation of uv1 cells is sufficient to induce Ca^2+^ activity, we used a pulled glass capillary probe to deliver a mechanical stimulus to the uv1 cells in intact animals (Fig. 3A). We used a motorized stage to move the worm body 50 μm towards the probe for 1 s. Direct prodding of uv1 triggered a strong and reproducible Ca^2+^ transient (Fig. 3B-C), similar to that seen during egg-laying events in freely behaving animals and following vulval muscle optogenetic stimulation. Direct stimulation of one uv1 cell triggers Ca^2+^ activity in the paired uv1 cell as well with no significant difference of Ca^2+^ transient amplitude (Fig. 3B and data not shown). In freely behaving worms, time from egg release to uv1 Ca^2+^ transient peak takes ∼3.2 s compared to that seen following prodding (5.6 s) or following optogenetic triggering (5.7s) (data not shown). To optimize the direct stimulation approach, we varied the displacement and stimulus duration. There was no detectable uv1 Ca^2+^ response at 10 μm displacement while we found nearly 100% of prodded uv1 cells showed significant Ca^2+^ responses upon 40 μm displacement (Fig. 3D). While only half of uv1s showed Ca^2+^ response following 20 μm displacement, when a Ca^2+^ transient was observed, the amplitudes were equivalent (Fig. 3E). Changes in stimulus duration also did not significantly affect whether the uv1 cells showed a Ca^2+^ transient with 0.5, 1 and 2 s showing equivalent responses (Fig. 3E). Interestingly, a second uv1 stimulation induced a significantly weaker Ca^2+^ response (Fig. 3C and 3F). This was not merely a consequence of delay on the stage, as animals waiting 2 min before stimulation showed similar uv1 Ca^2+^ transient amplitudes compared to those animals that were stimulated immediately (Fig. 3F). This suggests the uv1 cells lose their sensitivity to subsequent mechanical stimulations. Together, these data show the uv1 Ca^2+^ transients to prodding are all-or-none: small displacements fail to activate uv1 cells to a detectable degree while sufficiently large displacements trigger similar activity.

**Figure 3:**
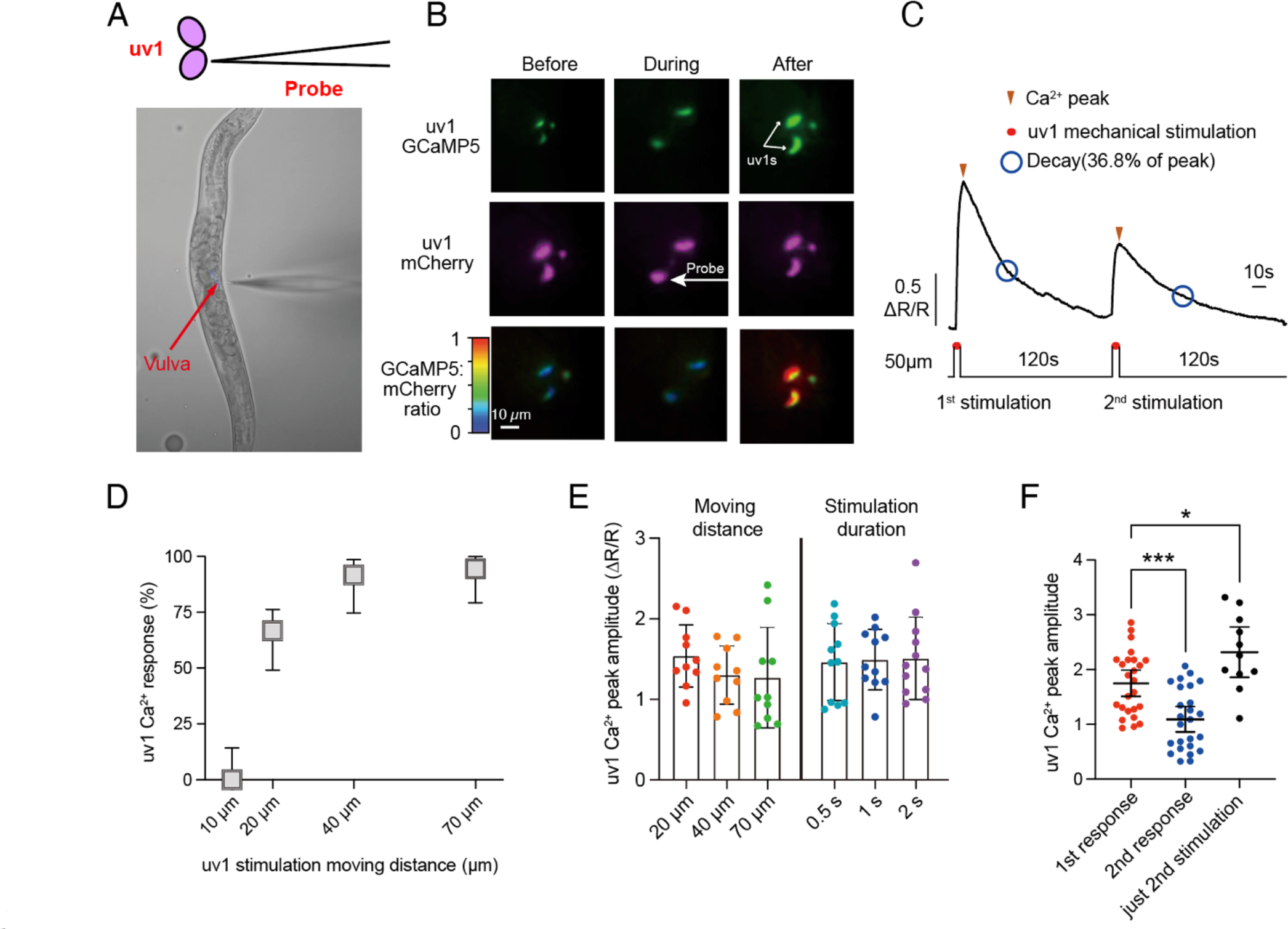
uv1 cells are strongly activated in response to manipulated direct mechanical stimulation. **(A)** Cartoon (top) of uv1 cells and micrograph demonstrating the mechanical prodding technique. A glass probe (arrowhead, black outline) is used to stimulate uv1 cells (purple). Brightfield micrograph (bottom) of the direct prodding method. Red arrow indicates the vulval position. The probe on the right points to the position of uv1 cells. The stage moves toward the needle, delivering the mechanical stimulus. **(B)** Micrographs of uv1 GCaMP5 and mCherry fluorescence and the GCaMP5/mCherry ratio before, during, and after direct prodding to uv1 cells. Arrowheads show uv1 position and induced GCaMP5 fluorescence following stimulation. **(C)** Averaged GCaMP5/mCherry ratio trace of uv1 Ca^2+^ transient activity in response to mechanical stimulation (n=20 animals). Red dots indicate prodding which lasts for 1 s. The stage displacement moves the worm 50 μm toward the probe which is placed opposed to the uv1 cells. A second stimulation occurs 2 minutes after first prodding event. Arrowheads indicate the Ca^2+^ transient peak. Blue open circles indicate the time of decay from Ca^2+^ transient peak down to 36.8%. **(D)** Box and whisker plot showing the percentage of uv1 cells showing Ca^2+^ transient activity relative to stage moving distance. Error bars represent ±95% confidence interval for the mean proportion. **(E)** Scatterplot of uv1 Ca^2+^ transient amplitude at different stage displacements (left) and stimulus durations (right). Error bars represent ±95% confidence interval for the mean. There are no significant differences between any bars. **(F)** Scatterplot of uv1s Ca^2+^ transient amplitude after first (red) and second (blue) stimulation and when only stimulated animal once after 120 s. Error bars represent ±95% confidence interval for the mean. A second stimulation induced a significantly weaker Ca^2+^ response compared to the first stimulation (p=0.0004, one-way ANOVA with Bonferroni’s correction for multiple comparisons). uv1 Ca^2+^ transient amplitudes in animals only receiving a single stimulus after 120 s are slightly increased relative to animals stimulated immediately (p=0.0203, one-way ANOVA with Bonferroni’s correction for multiple comparisons).

### uv1 activation is independent of synaptic transmission

Even though the timing of uv1 Ca^2+^ activity following stimulation suggested the uv1 cells responded directly to mechanical input, it remained possible that signals from other cells contribute to the response. We first tested the role of the HSN and/or VC neurons (Fig. 4A). We prodded the uv1 cells in animals bearing the *egl-1(n986dm)* mutation that causes the premature death the HSN command neurons (Conradt and Horvitz, 1999) or animals transgenically expressing Tetanus Toxin in the VC neurons (Kopchock et al., 2021). We found mechanical stimulation of uv1 cells in these animals triggered strong uv1 Ca^2+^ transients like wild type (Fig 4.B-C). Loss of HSNs, but not transmission from VCs, led to a modest but significant decrease in uv1 Ca^2+^ transient amplitude (Fig 4F), there was no difference in peak onset or decay time (data not shown). Overall, these results show direct mechanical stimulation activates uv1 without HSN or VC input.

**Figure 4:**
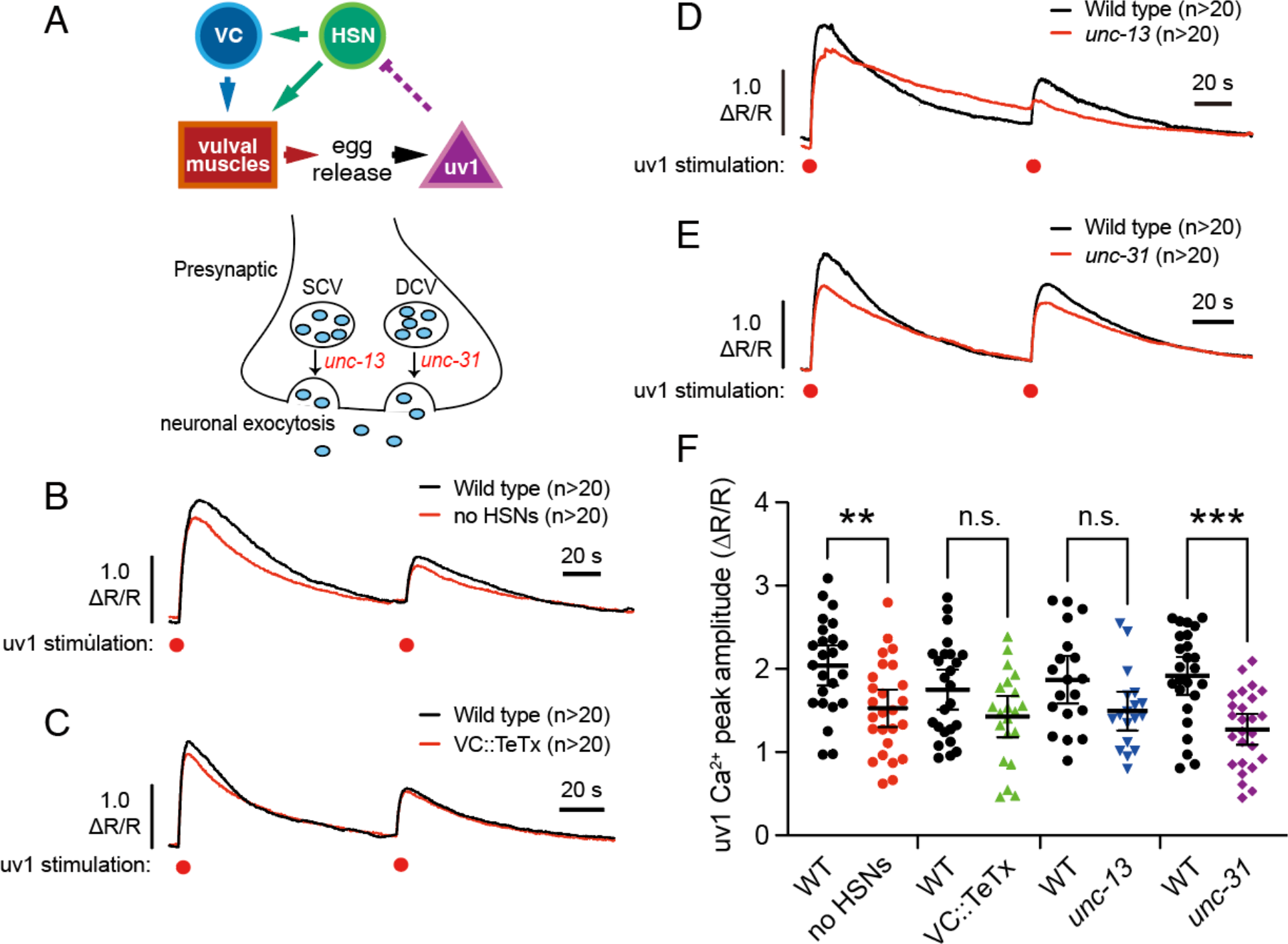
uv1 activation is independent of synaptic transmission. **(A)** Cartoon of the egg-laying circuit in *C. elegans* (top) and presynaptic neuronal exocytosis (bottom) **(B-E)** Averaged prodding-induced uv1 Ca^2+^ transient responses in wild-type, *egl-1(n986dm)* mutant animals **(B),** transgenic animals expressing Tetanus Toxin in the VC neurons **(C)** *unc-13(s69)* **(D)**, or *unc-31(e928)* **(E)** animals. n indicates number of animals tested and pooled (>20 per mutant with age-matched wild-type animals tested on same day). **(F)** Scatterplot showing uv1 Ca^2+^ transient amplitudes. ** represents p<0.01; *** represents p<0.001; n.s. represents p>0.05 (one-way ANOVA with Bonferroni’s correction for multiple comparisons).

To determine if uv1 activity in response to mechanical stimulation requires cells outside egg-laying circuit, we prodded *unc-13(s69)* mutants defective in synaptic vesicle exocytosis (Richmond et al., 1999) and *unc-31(e928)* CAPS mutants defective in dense core vesicle exocytosis (Speese et al., 2007). Both mutants showed similarly strong uv1 Ca^2+^ activity after mechanical stimulation compared to wild type worms (Fig 4D-E). Notably, uv1 Ca^2+^ transient amplitudes were significantly reduced in *unc-31(e928)* mutants (Fig. 4F), like that seen in *egl-1(n986dm)* mutants lacking HSNs. These results show that neurotransmitter or neuropeptide signaling is not strictly required for prodding-induced uv1 Ca^2+^ transient activity, although neuropeptide signaling, possibly from HSNs (Brewer et al., 2019), may modulate uv1 mechanosensory responses.

### uv1 mechanical activation can proceed without vulval muscle activity and contractility

Coordinated contraction of the vulval muscles opens the vulva for egg release (Brewer et al., 2019; Li et al., 2013; Olson et al., 2023). We have previously shown that the vulval muscles themselves respond to direct mechanical stimulation (Medrano and Collins, 2023), raising the possibility that vulval prodding stimulates vulval muscle activity and/or contractility that subsequently activates the uv1 cells. To test this, we prodded *unc-54(e190)* muscle myosin mutants with defective muscle contractility and egg laying (Moerman et al., 1982), but with strong injection-induced vulval muscle Ca^2+^ activity (Medrano and Collins, 2023). Prodding triggered strong uv1 Ca^2+^ transients in *unc-54(e190)* mutants (Fig. 5A), showing muscle contractility is not required. To test the role of vulval muscle electrical excitability, we transgenically expressed the histamine-gated Cl^-^ channel into the vulval muscles, allowing for potent silencing and egg-laying blockade following acute treatment with histamine (Ravi et al., 2018a). Prodding vulval muscle-silenced animals had no effect on prodding-induced uv1 Ca^2+^ activity (Fig. 5B). To separately test how vulval muscle electrical silencing affects uv1 mechanical activation, we transgenically expressed the A331T gain-of-function ERG channel mutant which strongly inhibits vulval muscle excitability and egg laying (Collins and Koelle, 2013). Prodding of uv1 cells in ERG gain-of-function mutants still triggered robust uv1 Ca^2+^ activity, although it was significant reduced in amplitude (∼35%; Fig. 5C, 5E), resembling results seen in animals lacking HSNs or CAPS neuropeptide signaling. ERG A331T is driven from a single-copy MosSCI transgene (Dhakal et al., 2022) using the vulval muscle-expressed *unc-103e* promoter/enhancer (Reiner et al., 2006), but we cannot exclude effects from expression in other cells. To eliminate the vulval muscles entirely, we prodded uv1 cells in *egl-17(n1377)* mutants with defects in sex myoblast migration (Burdine et al., 1997). We focused our prodding on the anterior uv1 cells since we previously showed that over 90% of *egl-17(n1377)* mutant animals lack anterior vulval muscle development and/or differentiation (Medrano and Collins, 2023). Direct prodding in these animals still triggered a strong uv1 Ca^2+^ transients (Fig. 5D) showing the vulval muscles are not required for uv1 mechanical activation. Together, our data show that vulval muscle activity and contractility are not required for the uv1 mechanical response to direct stimulation.

**Figure 5:**
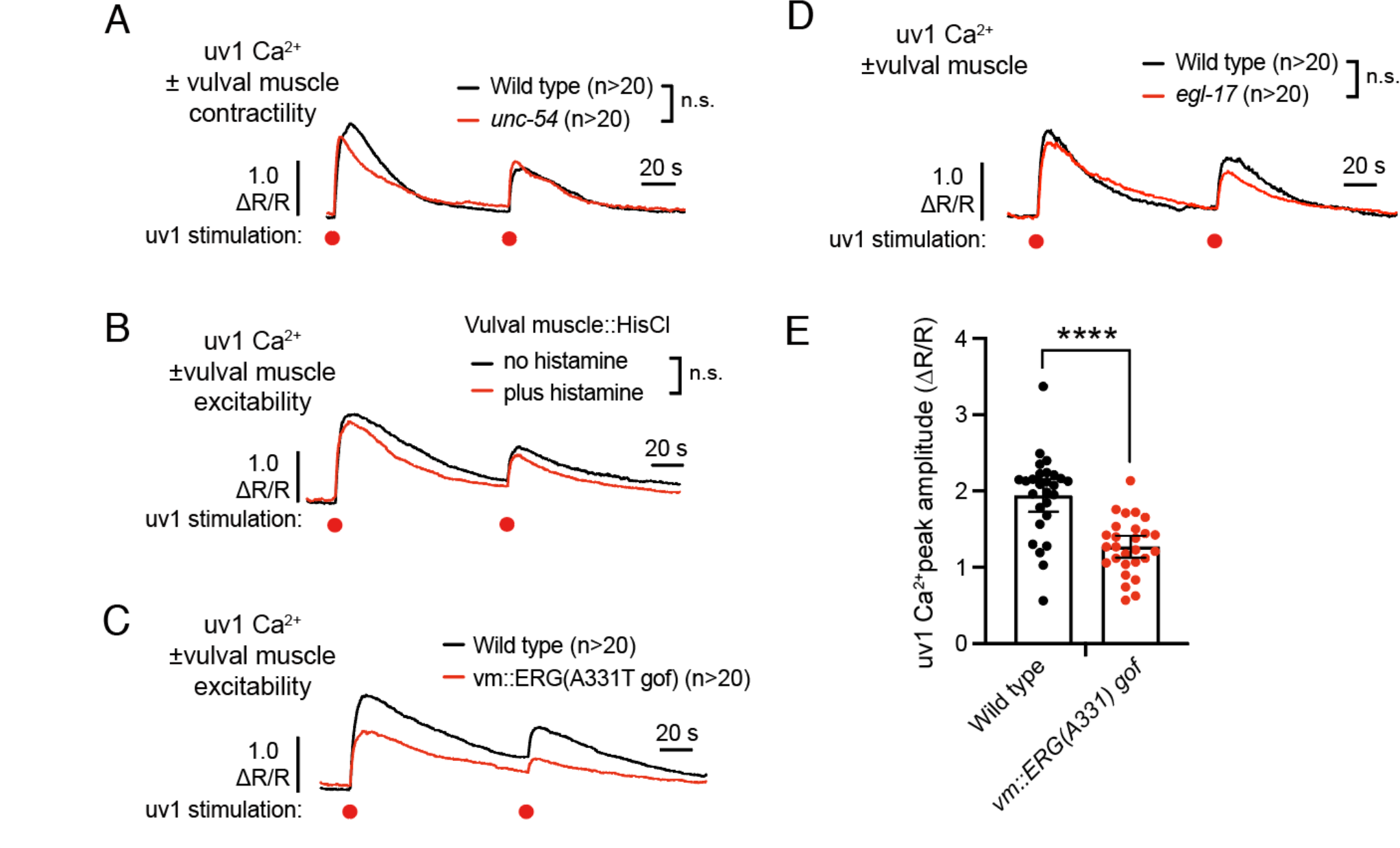
uv1 activation is independent of the vulval muscles. **(A-D)** Averaged prodding-induced uv1s Ca^2+^ responses of wild-type, *unc-54(e190)* **(A),** VM::HisCl worms with (red) or without (black) Histamine on the plates **(B)**, *unc-103(A331T)* gain of function mutants **(C)**, or *egl-17(n1377)* mutants lacking vulval muscle development / differentation **(D)**. **(E)** Scatterplot of uv1 Ca^2+^ transient peaks after direct stimulation in wild type (black) and *unc-103(A331T)* gain-of-function mutants (red). **** indicates p<0.0001 (Mann-Whitney test); n indicates number of animals tested per genotype or condition.

### uv1 mechanical activation does not require Piezo or TRPV channels

Piezo channels are mechanically gated and involved in many physiological processes in mammals (Coste et al., 2010). PEZO-1 is the sole ortholog of the Piezo protein in *C. elegans* and is expressed in vulval cells (Bai et al., 2020). *pezo-1(av149)* knockout mutants have deficiencies in brood size and ovulation (Bai et al., 2020), suggesting they may be involved in reproductive behaviors like egg laying. Previous patch clamp experiments from embryonic *C. elegans* cells have shown mechanosensitive PEZO-1 currents (Millet et al., 2022). To test if Piezo channels mediate uv1 mechanical activation, we prodded *pezo-1(av149)* mutants and measured uv1 Ca^2+^ activity. Prodding of *pezo-1(av149)* mutants still show strong uv1 Ca^2+^ transients (Fig. 6A) with no significant differences in Ca^2+^ transient kinetics or amplitude (Fig. 6E), indicating that Piezo channels are not required for the uv1 prodding response.

**Figure 6:**
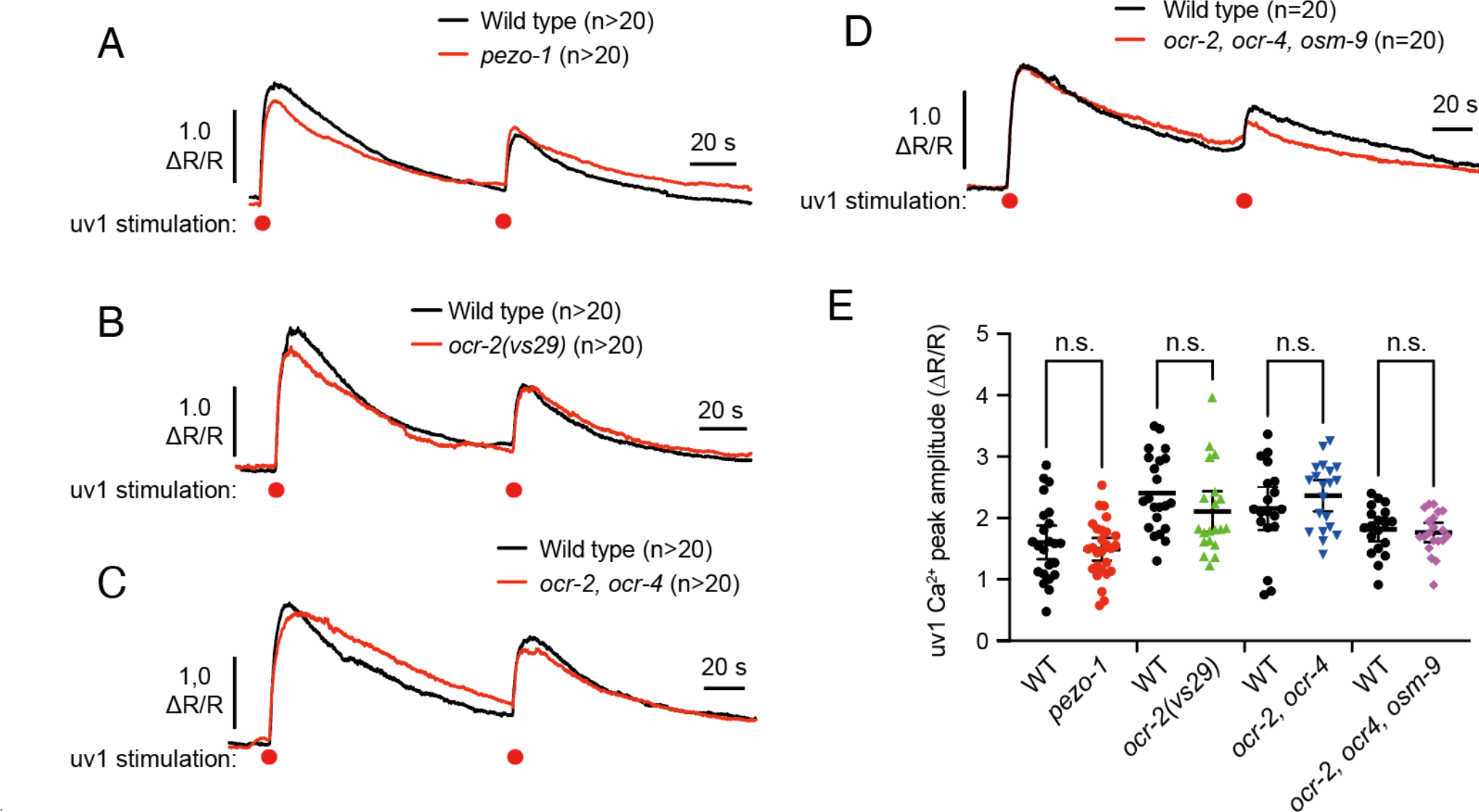
uv1 activation is independent of known mechanosensitive channels. **(A-D)** Averaged prodding-induced uv1s Ca^2+^ responses of wild-type (black); *pezo-1* **(A)** *ocr-2(vs29)* TRPV mutant **(B)**; *ocr-2(ak47), ocr-4(vs137)* TRPV double mutants **(C)**, or *ocr-2(ak47), ocr-4(vs137), osm-9(cx10) TRPV triple mutants* **(D)**. **(E)** Scatterplot showing uv1 Ca^2+^ transient peaks after direct stimulation. n.s. indicates p>0.05 (one-way ANOVA with Bonferroni’s correction for multiple comparisons). n indicates number of animals tested alongside age-matched wild-type control animals.

uv1 expresses other channels including TRPV channel that have been shown to facilitate mechanosensory responses (Birder et al., 2002). Prodding of *ocr-2(vs29)* mutant animals which show strongly hyperactive egg-laying behavior consistent with altered uv1 function showed similarly strong uv1 Ca^2+^ transient activity compared to wild-type control animals (Fig. 6B and 6E). Prodding of *ocr-2(ak47), ocr-4(vs137)* TRPV double mutants or *ocr-2(ak47), ocr-4(vs137), osm-9(cx10)* triple mutants triggered strong uv1 Ca^2+^ transients (Fig 6D-E). We found no differences in uv1 Ca^2+^ peak amplitude (Fig. 6E) or the timing of peak onset or decay time (data not shown). These data indicate that TRPV mechanosensitive channels are not required for uv1 mechanical activation.

### uv1 activation requires the EGL-19 L-type Ca^2+^ channel

L-type voltage-gated Ca^2+^ channels mediate Ca^2+^ entry and all-or-none action potential membrane depolarizations in *C. elegans* neurons and muscle cells (Gao and Zhen, 2011; Jospin et al., 2002; Liu et al., 2018). *C. elegans* has a single gene, *egl-19*, which encodes L-type Ca^2+^ channels (Lee et al., 1997). To test whether L-type Ca^2+^ channels are required for uv1 mechanical activation, we tested uv1 Ca^2+^ responses in animals where channel function was altered pharmacologically or genetically through loss- and gain-of-function mutations. Prodding *egl-19(n582)* mutants with reduced L-type Ca^2+^ channel function (Jospin et al., 2002; Lee et al., 1997) triggered uv1 Ca^2+^ transients with significantly reduced peak amplitude compared to wild-type worms (Fig. 7B). Conversely, prodding of *egl-19(n2368)* gain-of-function mutant animals in which L-type Ca^2+^ channel activity is increased (Laine et al., 2014) showed similarly strong uv1 Ca^2+^ transients (Fig. 7C), and these transients decayed more slowly, increasing from 43 s in wildtype to 74 s in *egl-19(n2368(gf))* mutant animals (Fig. 7D). These results show that EGL-19 L-type voltage-gated Ca^2+^ channels promote uv1 Ca^2+^ activity in response to direct mechanical stimulation.

**Figure 7:**
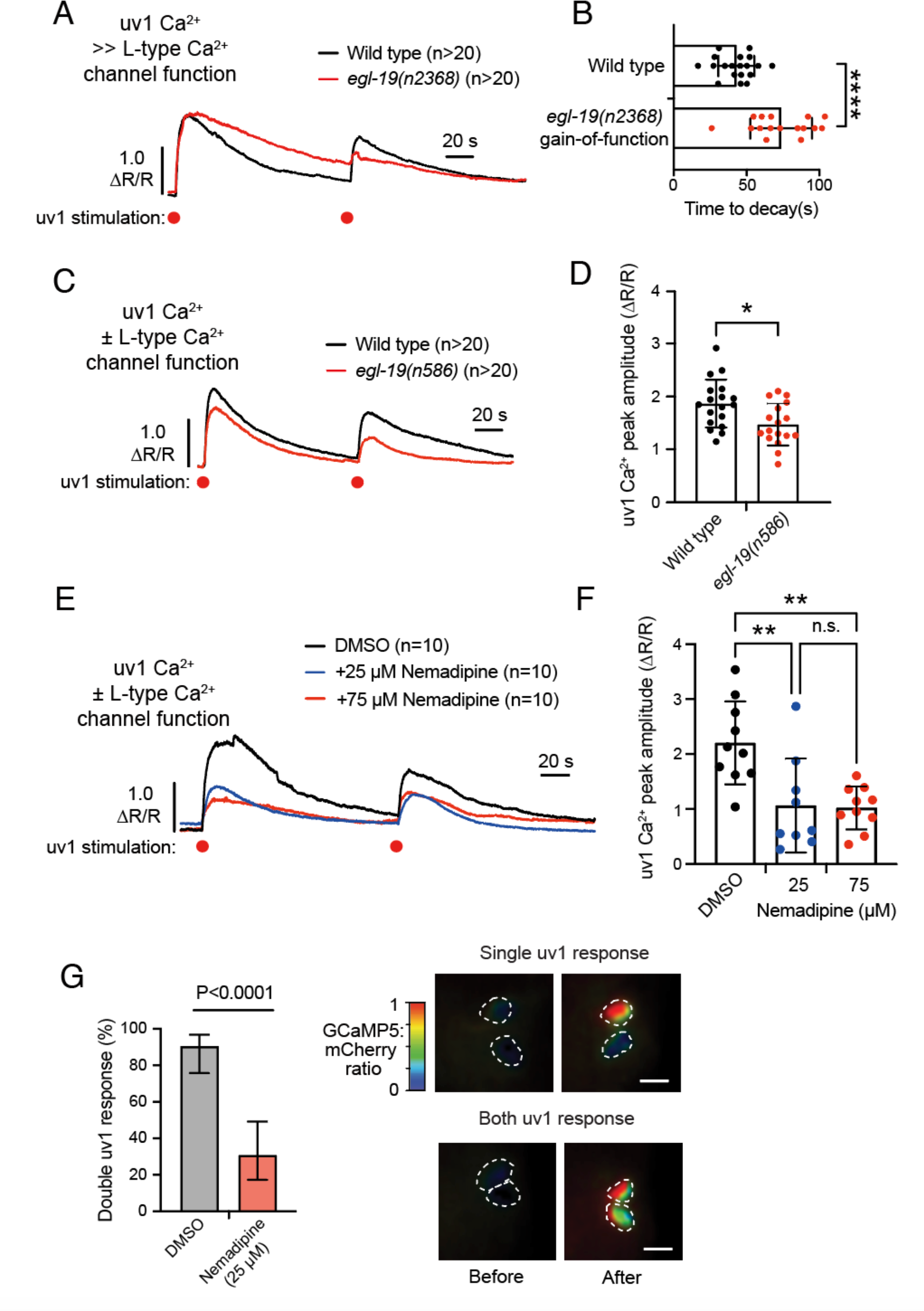
uv1 activation requires EGL-19 voltage-gated Ca^2+^ channels. **(A)** Averaged prodding-induced uv1s Ca^2+^ responses of wild-type (black) and *egl-19(n2368)* gain-of-function mutant animals (red). **(B)** Scatterplot of Ca^2+^ transient decay times. Bar indicates mean ±95% confidence interval, circles indicate prodding responses (n>20 animals). **** represents p<0.0001 (Mann-Whitney test). **(C)** Averaged prodding-induced uv1s Ca^2+^ responses of wild-type (black) and *egl-19(n586)* loss-of-function mutants (red). **(D)** Scatterplot of Ca^2+^ transient amplitudes. Bar indicates mean ±95% confidence interval; circles indicate prodding responses (n>20 animals); * indicates p<0.05 (Mann-Whitney test). **(E)** Averaged prodding-induced uv1 Ca^2+^ responses of wild-type worms treated with 0.25% DMSO (black), 25 μM nemadipine (blue) or 75 μM nemadipine (red). **(F)** Scatterplot showing uv1 Ca^2+^ transient amplitudes. Bar indicates mean ±95% confidence interval; circles indicate prodding responses (n>20 animals); ** represents p<0.01. n.s. represents p>0.05 (one-way ANOVA with Bonferroni’s correction for multiple comparisons). **(G)** Left, bar graphs (±95% confidence interval for the proportion) comparing percentage of animals showing Ca^2+^ transient responses in both anterior and posterior uv1 cells following treatment with 0.25% DMSO (black) or 25 μM nemadipine (red); p<0.0001 (Fischer’s exact test). Right, micrographs showing single and both uv1 response before and after direct prodding. Scale bar =10 μm.

Since *egl-19* null mutations are lethal (Jospin et al., 2002), we used the specific chemical inhibitor nemadipine to block channel function acutely in otherwise wild-type animals (Kwok et al., 2006). Animals treated with 25 or 75 μM nemadipine showed significantly reduced uv1 Ca^2+^ transient amplitudes following direct prodding compared to 0.25% DMSO control animals (Fig. 7E). We found no significant differences between 25 μM and 75 μM nemadipine treatments (Fig. 7F), suggesting the remaining uv1 Ca^2+^ activity is not caused by insufficient channel inhibition. Subsequent experiments used 25 μM nemadipine treatment in wild-type worms with a higher sample size to further determine the role of L-type Ca^2+^ channels in in uv1 mechanical activation. Treatment with 25 μM nemadipine caused a 50% reduction in uv1 Ca^2+^ transient amplitude following prodding (Fig. 7F). We also found nemadipine reduced the coordinated uv1 response following prodding. In untreated animals, 90% of uv1 cells on both the anterior and posterior side of the vulva show activation, no matter which side of uv1 cells were initially prodded. This coordinated uv1 Ca^2+^ activity was reduced to 30% following treatment with 25 μM nemadipine (Fig. 7G). The reduction in uv1 Ca^2+^ transient amplitude observed was not a result of impaired coordination, as nemadpine reduced uv1 Ca^2+^ transient responses even in those uv1 cells that showed a Ca^2+^ response when they were measured separately. Nemadipine treatment reduced, but did not eliminate, the observed uv1 Ca^2+^ transient peak amplitude compared DMSO-treated control animals, suggesting EGL-19 L-type Ca^2+^ channels facilitate the full Ca^2+^ transient response following uv1 mechanical activation.

### uv1 activation is independent of UNC-2 P/Q/N-type Ca^2+^ channel and UNC-68 ryanodine receptor

*C. elegans* expresses other voltage-gated channels (Bargmann, 1998). *unc-2* encodes the sole P/Q/N-type voltage-gated calcium channel (Mathews et al., 2003) which is expressed in the HSN and VC neurons (Mathews et al., 2003). *unc-2(e55)* mutants show hyperactive egg-laying behavior (Schafer and Kenyon, 1995) and altered egg laying in response to serotonin (Schafer et al., 1996). Prodding and uv1 Ca^2+^ imaging in *unc-2(e55)* mutant showed strong Ca^2+^ transients (data not shown). In *C.elegans*, L-type (EGL-19) and P/Q/N-type (UNC-2) channels localize differentially in synapses and mediate distinct pools of vesicle release (Mueller et al., 2023). To test if EGL-19 and UNC-2 channel function together in uv1 mechanical activation, we treated *unc-2(e55)* mutants with 25 μM nemadipine and performed direct prodding. Loss of both UNC-2 and EGL-19 channel function failed to reduce uv1 Ca^2+^ activity further (Fig. 8A, 8B), suggesting these two channels do not act in parallel to facilitate uv1 Ca^2+^ activity following mechanical stimulation.

*C. elegans* also expresses ryanodine receptors (RyRs) which belong to family of intracellular (ER) calcium channels (Maryon et al., 1996). A previous study showed that UNC-68 RyRs enhance L-type Ca^2+^ channel (EGL-19)-mediated synaptic vesicle release (Mueller et al., 2023). To test if RyR acts in parallel to EGL-19 in uv1 Ca^2+^ transient responses following mechanical activation, we performed prodding in *unc-68(e540)* mutants following treatment with 25 μM nemadipine. Prodding and uv1 Ca^2+^ imaging in *unc-68(e540)* mutant still showed strong Ca^2+^ transients with a slight decrease in transient decay time (P=0.037) (data not shown), consistent with RyR channels sustaining uv1 Ca^2+^ responses following stimulation. However, loss of both UNC-68 and EGL-19 channel function did not eliminate the residual uv1 Ca^2+^ activity following stimulation (Fig 8C, 8D), showing that ryanodine receptor UNC-68 RyRs are not required for uv1 mechanical activation in the absence of L-type Ca^2+^ channel function.

## Discussion

The mechanical activation of neuroendocrine cells reveals the intricate interplay between physical stimuli and cellular signaling mechanisms. In this study, we investigated how the uv1 neuroendocrine cells of the *C. elegans* egg-laying behavior circuit are activated. Our findings demonstrated the uv1 cells are mechanically activated by vulval muscle contraction. Importantly, our data revealed that the mechanical activation of uv1 cells occurs independently of egg release, suggesting the uv1 cells respond to and report instances of vulval opening. To test whether uv1 mechanical activation is direct, we devised a prodding assay where the uv1 cells are stimulated with glass probes. We found that mechanical stimulation triggers a strong and “all-or-none” uv1 Ca^2+^ transient. Notably, uv1 cells respond most robustly to the first stimulation with subsequent mechanical stimulations eliciting weaker Ca^2+^ transients. Using mutants with specific defects in egg-laying circuit activity, development, or neurotransmission our experiments revealed uv1 mechanical activation can be direct. While uv1 Ca^2+^ transients were reduced in mutants lacking the HSNs, the underlying prodding response was still observed. This resembles our results with *unc-31(e928)* CAPS mutants, suggesting neuropeptide transmission from cells including HSN promotes the uv1 response. HSNs release serotonin and NLP-3 neuropeptides to stimulate vulval muscle excitability (Brewer et al., 2019; Shyn et al., 2003; Waggoner et al., 1998), possibly by activation of G protein coupled receptors to modulate activity of postsynaptic L-type Ca^2+^ channels and/or voltage-gated K^+^ channels (Lee et al., 1997; Waggoner et al., 1998). As retrograde feedback of egg accumulation and vulval muscle Ca^2+^ activity facilitates burst-firing in HSNs (Medrano and Collins, 2023; Ravi et al., 2018a; Ravi et al., 2021), defects in HSN transmission may cause ancillary effects on uv1 mechanical activation and release of tyramine that signals to terminate the egg-laying behavior state (Collins et al., 2016).

**Figure 8:**
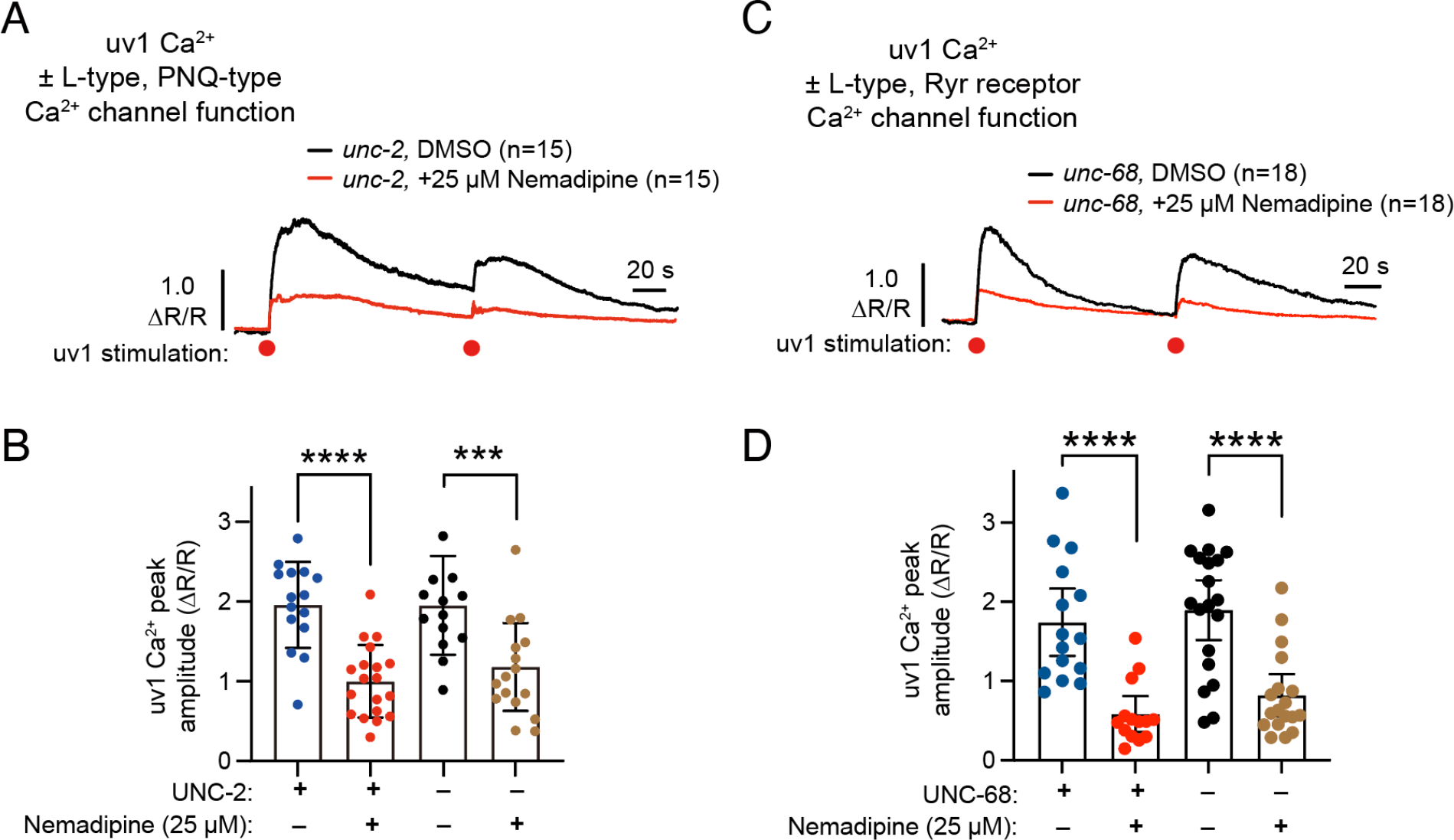
Other Ca^2+^ channels do not act redundantly with L-type Ca^2+^ channels in uv1 mechanical activation. **(A)** Averaged prodding-induced uv1 Ca2+ responses in *unc-2(e55)* mutant animals following treatment with 0.25% DMSO (black) or 25 μM nemadipine (red) (n=15 animals). **(B)** Scatterplot showing uv1 Ca^2+^ transient peak amplitudes after direct stimulation in wild-type and *unc-2(e55)* mutants following treatment with 0.25% DMSO or 25μM nemadipine. Bar indicates mean ±95% confidence interval; **** indicates p<0.0001; *** indicates p-value<0.001 (one-way ANOVA with Bonferroni’s correction for multiple comparisons.) **(C)** Averaged prodding induced uv1 Ca2+ responses in *unc-68(e540)* mutant animals following treatment with 0.25% DMSO (black) or 25 μM nemadipine (red) (n=18). **(D)** Scatterplot showing showing uv1 Ca^2+^ transient peaks after direct stimulation in wild-type or *unc-68(e540)* mutant animals following treatment with 0.25% DMSO or 25 μM nemadipine. Bar indicates mean ±95% confidence interval; **** indicates p<0.0001 (one-way ANOVA with Bonferroni’s correction for multiple comparisons).

How is uv1 mechanically activated? Previous studies showed that TRPV and Piezo mechanosensitive channels are expressed in the uv1 cells and vulva (Bai et al., 2020; Jose et al., 2007) suggesting these channels as candidates (O’Neil and Heller, 2005). However, we find that mutants animals lacking three uv1-expressed TRPV channels or Piezo still showed strong uv1 Ca^2+^ transients during egg laying and in response to direct prodding, suggesting TRPV or Piezo channels do not mediate uv1 mechanical activation. This suggests either functional redundancy among these channels and/or the presence of additional mechanosensitive channels in uv1 cells that detect physical stimuli and trigger uv1 cellular activation. In other *C. elegans* mechanosensory neurons, previous work has shown that DEG/ENAC channel MEC-4 complex is essential for transducing external mechanical force into ionic currents in touch neurons (O’Hagan et al., 2005). TRPA1 encodes for an ion channel serving as sensor molecule in enterochromaffin neuroendocrine cells may regulating gastrointestinal function in mammals (Nozawa et al., 2009). Its ortholog TRPA-1 is also activated by mechanical stimulation and contributes to mechanosensory behaviors such as nose-touch response in *C. elegans* (Kindt et al., 2007). As uv1 is not well-represented in community-derived, cell-specific RNAseq resources, further work will be required to identify and test the contribution of these and other candidate channels.

Our work shows an important role for the EGL-19 L-type voltage-gated Ca^2+^ channel. Mutations that increase EGL-19 channel function sustain uv1 Ca^2+^ activity after stimulation while chemical or genetic inhibition of EGL-19 channel function significantly impair it. Surprisingly, UNC-68 ryanodine receptors, responsible for intracellular Ca^2+^ release and essential for EGL-19-mediated synaptic transmission in *C. elegans* (Lanner et al., 2010; Mueller et al., 2023), were not involved in uv1 mechanical activation. Furthermore, our findings revealed that the P/Q/N type Ca^2+^ channel (Mathews et al., 2003) does not cooperate alongside EGL-19 in mediating uv1 mechanical activation. At present, the source of the Ca^2+^ transients we observe is not clear, although our work showing a critical role for L-type Ca^2+^ channels is consistent with a significant fraction of the observed Ca^2+^ transients coming from an extracellular source. Future work will be required to determine whether the remaining Ca^2+^ signals we observe arises from external sources such as mechanically activated channel(s) or internal sources (or both). Previous study showed that voltage-gated Ca^2+^ channels can amplify mechanosensory signals and increase mechanoreceptors sensitivity in touch neurons of mice (Wang and Lewin, 2011). Given our data that uv1 mechanical response does not require neurotransmission from other cells in egg-laying circuit, EGL-19 may similarly act as amplifier of mechanosensory signals. We postulate that unidentified mechanical receptors in uv1 cells that perceive physical stimuli and subsequently signal to the voltage-gated Ca^2+^ channel EGL-19, initiating uv1 cellular activity.

Our findings further suggest that most or all cells of the *C. elegans* behavior circuit are mechanically responsive. The vulval muscles are regulated by both internal mechanosensory feedback of egg accumulation and external mechanical stimulation (Medrano and Collins, 2023). Feedback of vulval muscle Ca^2+^ activity drives retrograde feedback to the presynaptic HSN neurons (Ravi et al., 2018a), allowing for proper stretch-dependent modulation of HSN activity and the egg-laying active state in animals with sufficient eggs to lay (Ravi et al., 2021). We have also shown that the VC neurons show peak activity coincident with vulval muscle Ca^2+^ activity and contractility (Collins et al., 2016). Direct optogenetic stimulation of the vulval muscles similarly triggers VC Ca^2+^ activity which is reduced in *unc-54* muscle myosin mutants (Kopchock et al., 2021). Processes from the VC neurons extend ventrally along the vulval opening and dorsally where they make synapses onto the vulval muscles near the HSNs. The uv1 cells are situated even more laterally, just dorsal to the VC and HSN synapses (Ravi et al., 2018a), extending processes down to the VulF vulval epithelial cells and dorsally to the utse uterine seam cell (Newman et al., 1996). Based on this anatomy, we predict VC mechanically responds to smaller scale vulval openings while uv1 only responds when the vulva opens completely, deforming uv1 cellular connections. This model mirrors results seen in vertebrates, where different touch receptor neurons detect different kinds of mechanical stimuli including touch and pressure, contributing to the perception of tactile sensations while maintaining sensitivity to stimulus (Geffeney and Goodman, 2012). The cells in the egg-laying circuit may be providing both functions: sensing acute changes in egg accumulation, releasing serotonin and neuropeptides to drive sufficient vulval contraction for egg release, but not allowing too much activity that animals are unable to maintain vulval integrity (Gianakas et al., 2023).

We also find that EGL-19 channels facilitate the coordination of uv1 Ca^2+^ activity across the vulva. We consistently observe synchronized Ca^2+^ activity in both anterior and posterior uv1 cells in response to both vulval opening and direct prodding stimuli. Notably, blocking L-type Ca^2+^ channels led to Ca^2+^ transients only in the prodded uv1 with the opposite uv1 failing to show a robust Ca^2+^ transient. Such coordination may be mediated by adherens and/or gap junctions. uv1 extends processes that mediate contact between the utse uterine seam cell and the VulF cells of the vulval lumen (Newman et al., 1996). Such contacts may also be sites of gap junctions, connecting cells electrically and biochemically through diffusion of ions (such as Ca^2+^), second messengers, and small metabolites (Mese et al., 2007; Simonsen et al., 2014). Previous work showed that UNC-7 and UNC-9 innexin gap junctions are required for gap junction in electrically coupled networks (Walker and Schafer, 2020). Furthermore, UNC-7 can form hemichannels that drive gentle mechanosensation within touch neurons, as well as in harsh touch sensation within the PVD polymodal nociceptors (Walker and Schafer, 2020). These findings suggested innexins like UNC-7 and/or UNC-9 may be candidate uv1 mechanoreceptors. However, the direct prodding results of *unc-7* and *unc-9* mutants show normal uv1 prodding responses, indicating they are not strictly required (data not shown). In *C. elegans*, there are 25 identified innexin genes which are expressed widely in neurons or tissues (Simonsen et al., 2014). As nearly all cells in both the egg-laying circuit and the somatic gonad make gap junctions (White et al., 1986), we predict innexins coordinate cellular Ca^2+^ activity in uv1 and other cells during egg laying.

In conclusion, our study provides a comprehensive exploration of uv1 neuroendocrine mechanical activation, uncovering novel aspects of the regulatory mechanisms. Our work suggests vulval opening activates unidentified mechanoreceptors, facilitating subsequent activation of L-type Ca^2+^ channels. Future work determining what molecules facilitate cellular communication and synchronized cellular activity will benefit further understanding of cell coordination and innexins functions.

## MATERIALS AND METHODS

### Worm culture

*C. elegans* worms were maintained at 20°C on Nematode Growth Medium (NGM) agar plates with *E. coli* OP50 as a source of food as described previously (Brenner, 1974). All experiments involving adult animals were performed using adult hermaphrodites staged 24 h past the late L4 larval stage. A list of strains, mutants, and transgenes used in this study can be found in **Table 1**.

**Table 1.**
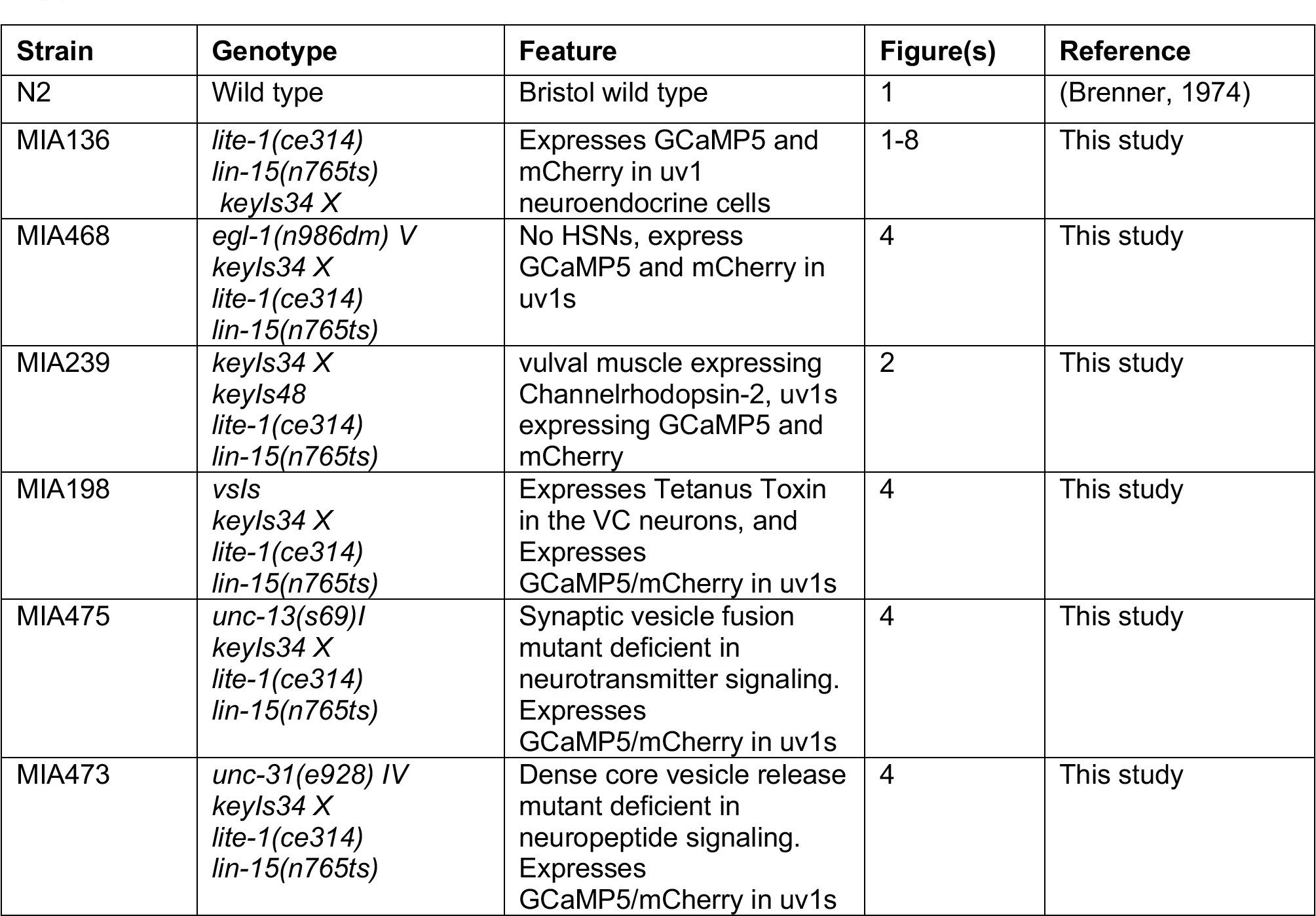
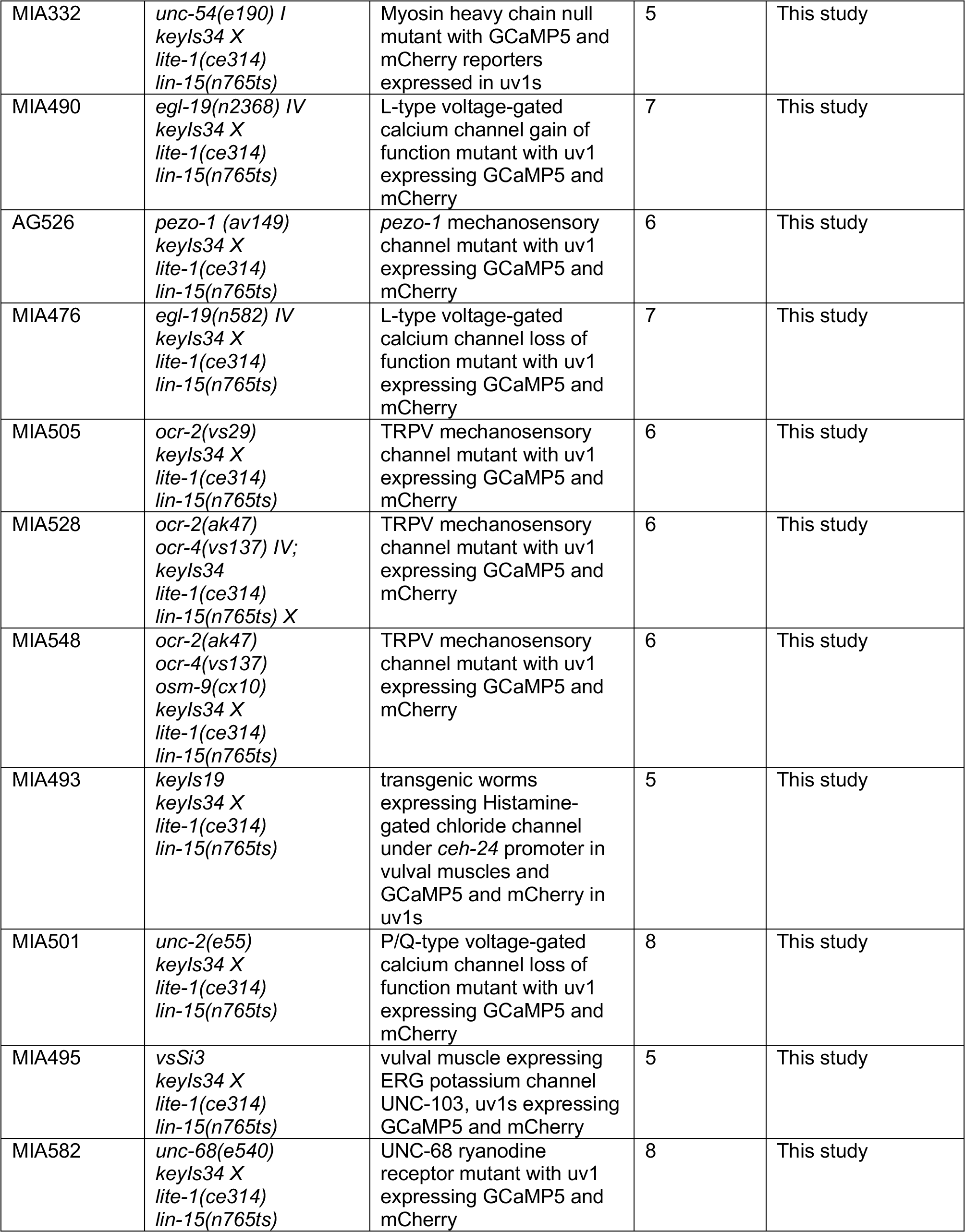
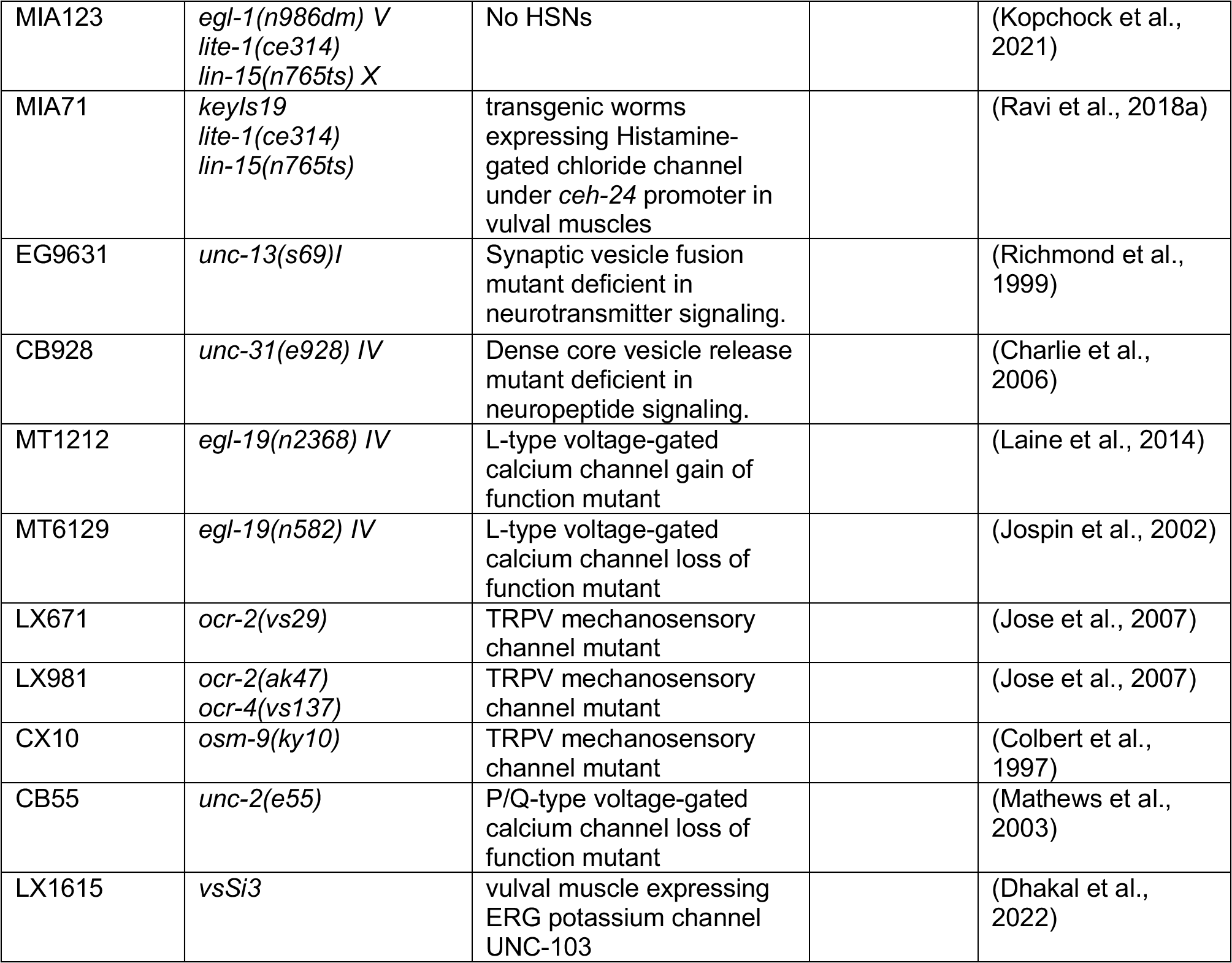

### Calcium reporter transgenes and strain construction

#### uv1 Ca^2+^ activity

Ca^2+^ activity of uv1 neuroendocrine cells was recorded in adult animals using MIA136 [*keyIs34 lite-1(ce314) lin-15(n765ts) X*] expressing GCaMP5 and mCherry from the *tdc-1* promoter. Oligos Ptdc-1new-fwd: GCG GCA TGC CAC CTA ACT TCG TCG GTC and Ptdc-1-new-rev: GCG CCC GGG GAT CCT TGG GCG GTC CTG were used to amplify *tdc-1* promoter from a ChR2 plasmid. The amplicons were digested with SphI and XmaI to generate a 1.5 kb PCR product which was then ligated into similarly digested pKMC281 (mCherry) and pKMC284 (GCaMP5) plasmids, generating pAB5 (*tdc-1p::mCherry*) and pAB6 (*tdc-1p::GCaMP5*). 20 µg/ml of pAB6 (*tdc-1p::GCaMP5*) and 5 µg/ml of pAB5 (*tdc-1p::mCherry*) along with 50 µg/ml of pLI5EK (*lin-15* rescue plasmid) were injected into LX1832 [*lite-1(ce314) lin-15(n765ts)* X] hermaphrodites. The extrachromosomal arrays were integrated using UV and trimethylpsoralen, and strains carrying an integrated version of the GCaMP5/mCherry transgene were then backcrossed six times to LX1832 animals to generate MIA136 [*keyIs34 lite-1(ce314) lin-15(n765ts) X*].

To image uv1 Ca^2+^ activity in HSN-deficient mutant, MIA468 [*egl-1(n986dm)* V; *keyIs34 lite-1(ce314) lin-15(n765ts)* X] was generated. MIA136 males were crossed with MIA123 [*egl-1(n986dm); lite-1(ce314), lin-15(n765ts)*] hermaphrodites lacking HSNs (Conradt and Horvitz, 1999). Fluorescent cross-progeny L4s were then cloned and allowed to self-fertilize. Egl, mCherry+ hermaphrodites were then selected, and the *egl-1(n986dm)* mutation was confirmed by genotyping using the oligos: 5’-CTC TGT TCC AGC TCA AAT TTC C-3’ (forward) and 5’-GTA GTT GAG GAT CTC GCT TCG GC-3’ (reverse) followed by NciI enzyme digestion.

To image uv1 Ca^2+^ activity in neurotransmission defective worms, MIA473 [*unc-31(e928) IV*; *keyIs34 lite-1(ce314) lin-15(n765ts)* X] and MIA475 [*unc-13(s69) I*; *keyIs34* X; *lite-1(ce314) lin-15(n765ts)* X] were generated. To generate these strains, MIA136 males were crossed with uncoordinated CB928 [*unc-31(e928) IV*] CAPS (Charlie et al., 2006) or EG9623 [*unc-13(s69) I*] (Richmond et al., 1999) hermaphrodites. Resulting mCherry+ hermaphrodites were picked and allowed to self-fertilize. Uncoordinated, mCherry+ F2 progeny were then and allowed to self-fertilize until both the uncoordinated and uv1 fluorescence were homozygous. *lite-1(ce314)* was confirmed to be homozygous by genotyping.

To image uv1 Ca^2+^ activity in L-type calcium-gated channel EGL-19 mutants, MIA476 [*egl-19(n582) IV*; *keyIs34 lite-1(ce314) lin-15(n765ts)* X] and MIA490 [*egl-19(n2368) IV*; *keyIs34 lite-1(ce314) lin-15(n765ts)* X] strains were generated. MIA136 [*keyIs34 lite-1(ce314) lin-15(n765ts) X*] males were crossed with MT6129 [*egl-19(n582) IV*] (Jospin et al., 2002) loss-of-function or MT1212 [*egl-19(n2368) IV*] gain-of-function mutant (Laine et al., 2014) hermaphrodites. Cross-progeny were selected and self-fertilized until uv1 fluorescence and *egl-19* defects were homozygous by phenotype. *lite-1(ce314)* was confirmed to be homozygous by genotyping.

To image uv1 Ca^2+^ activity after vulval muscle silencing, MIA493 [*keyIs19*; *keyIs34*; *lite-1(ce314) lin-15(n765ts)* X], MIA495 [*vsSi3*; *keyIs34*; *lite-1(ce314) lin-15(n765ts)* X] and MIA332 [*unc-54(e190), keyIs48*; *keyIs34 lite-1(ce314) lin-15(n765ts)* X] were generated. MIA136 males were crossed with MIA71 [*keyIs19; lite-1(ce314) X; lin-15(n765ts)X*] strain expressing HisCl in the vulval muscles (Ravi et al., 2018a). mCherry+ cross-progeny were selected and allowed to self-fertilize. F2 progeny were then put onto NGM plates containing 5 mM histamine for 2 h, and induced Egl worms were selected. Subsequent rounds of phenotyping confirmed strains that were homozygous for both uv1::GCaMP5 and vm::HisCl transgenes. To generate MIA495, MIA136 males were crossed with LX1615 [*vsSi3 II*; *lite-1(ce314) lin-15(n765ts)* X] (Dhakal et al., 2022) animals expressing the A331T gain-of-function ERG mutant that leads to vulval muscle silencing and inhibition of egg laying (Collins and Koelle, 2013; Dhakal et al., 2022). Resulting cross-progeny L4s were then picked, allowed to self-fertilize, and then homozygous mCherry+, Egl, *lite-1(ce314)* animals were picked and kept as MIA495. To generate MIA332, MIA298 [*unc-54 (e190) I; lite-1 (ce314), lin-15 (n765ts) X*] hermaphrodites were mated with MIA239 [*keyIs48*; *keyIs34*; *lite-1(ce314) lin-15(n765ts)* X] males. 6 non-Unc F1 L4 progeny with uv1 fluorescence were individually cloned and allowed to self. The presence of homozygous *unc-54* and uv1 GCaMP/mCherry was confirmed by the Unc phenotype and uv1 fluorescence.

To image uv1 Ca^2+^ activity in the *unc-2* loss of function mutant, MIA501 [*unc-2(e55)*; *lite-1(ce314) lin-15(n765ts)* X] was generated. MIA136 males were crossed with CB55 [*unc-2(e55)*] with hyperactive egg laying and uncoordinated locomotion (Mathews et al., 2003). Fluorescent cross-progeny L4’s was allowed to self-fertilize, and homozygous Unc, mCherry+, and *lite-1(ce314)* animals were kept.

To image uv1 Ca^2+^ activity in TRPV mechanosensory channel mutants, MIA505 [*ocr-2(vs29)*; *keyIs34 lite-1(ce314) lin-15(n765ts)* X], MIA528 [*ocr-2(ak47) ocr-4(vs137); keyIs34 lite-1(ce314) lin-15(n765ts)* X], MIA548 [*ocr-2(ak47) ocr-4(vs137) osm-9(ky10); keyIs34 lite-1(ce314) lin-15(n765ts)* X] were created. To generate these strains, MIA136 males were crossed with LX671 [*ocr-2(vs29)*] or LX981 [*ocr-2(ak47); ocr-4(vs137)*] (Jose et al., 2007) hermaphrodites. Resulting cross-progeny L4s were picked and allowed to self-fertilize. *ocr-2(vs29)* were confirmed to be homozygous through the true-breeding hyperactive egg-laying phenotype (Jose et al., 2007). Homozygous *ocr-2(ak47)* and *ocr-4(vs137), ocr-2(ak47)* mutant animals were identified through PCR genotyping using the following oligos: 5’-CGT TAA CAC TTG CGG CGA AAT TGG-3’ (ocr-2 forward), 5’-CCC GCC AAT ACG ATG GTT TGG-3’ (wild-type reverse), 5’-GCA GCA TTT AAC TGG ACC CCA ATT T-3’ (wild-type forward). *ocr-4(vs137)* was confirmed to be homozygous using oligos: 5’-GTG TTA CGA GAA GCA TAT AGA CTA CG-3’ (forward) and 5’-CCA AGC ATT ATG AGT AAC GTT GCG G-3’ (reverse), generating MIA528 [*ocr-2(ak47) ocr-4(vs137); keyIs34 lite-1(ce314) lin-15(n765ts)* X]. N2 males were then crossed with MIA528 hermaphrodites and the mCherry+, heterozygous TRPV double mutant cross-progeny were then mated with CX10 [*osm-9(ky10)*] hermaphrodites. mCherry+ cross-progeny L4s were picked singly onto plates and allowed to self. From these, 24 mCherry+ hermaphrodites were picked singly for genotyping for *ocr-2(ak47)* by PCR as described and *osm-9(ky10)* by Tetra-ARMS PCR using oligos: 5’-AGC TTA AAG TCT TCG GGT AGG AAG A-3’ (outside forward); 5’-CGA AAA TTT GAT CCA TCT TTT TGT TG-3’ (outside reverse); 5’-ATG GCT AGG TGG AGG GCT GAG TA-3’ (inside forward); 5’-CCT GAA ACA TTG TGT TAT CTT TAG TCC-3’ (inside reverse). *osm-9(ky10)* homozygous, *ocr-2(ak47)* heterozygous animals were selected and allowed to self-fertilize. From these, eight hermaphrodites were randomly selected and genotyped for both *ocr-2(ak47)* and *osm-9(ky10). osm-9(ky10)* homozygosity was then confirmed using Sanger sequencing while *ocr-4(vs137)* was confirmed by PCR-based genotyping as above.

### Ratiometric Ca^2+^ imaging and brightfield recording

Ratiometric Ca^2+^ imaging of uv1 activity of alive worms were performed as described (Ravi et al., 2018b). GCaMP5 and mCherry fluorescence was excited at 470 nm and 590 nm, respectively, using a Colibri.2 LED illumination system and channels were separated by a Hamamatsu W-View Gemini beam splitter. Ca^2+^ recordings were collected at 20 fps and a 256×256 pixel resolution (4x4 binning) using a Hamamatsu Flash 4.0 V2 sCMOS camera recording at 16-bit depth through a 20x Apochromatic objective (0.8 NA) on a Zeiss Axio Observer microscope. To record worm behaviors during ratiometric Ca^2+^ imaging, a Grasshopper 3 camera (FLIR) was used to capture 2 × 2 binned, 1,024 × 1,024 8-bit JPEG image sequences. For prodding experiments, each worm was recorded 20 s prior to first prodding to set up baseline levels for uv1 activity, and recordings were completed within 300 s to avoid effects of prolonged incubation in halocarbon oil (see below). For ChR2 experiments, each worm was recorded under the same condition for 120 s. For freely behaving animals, each worm was recorded 5 minutes after the first egg was laid. Worm recordings that did not have an egg-laying event in the first 30 minutes were excluded from the data analysis. Ratiometric Ca^2+^ recordings were then imported into Volocity (Version 6.5.1, Quorum Technologies, Inc.) for subsequent GCaMP5/mCherry ratiometric analysis. Ca^2+^ quantitation of GCaMP5/mCherry ratio changes (ΔR/R), Ca^2+^ transient peaks were analyzed through MATLAB as described (Collins et al., 2016; Ravi et al., 2018b).

### Prodding protocol

To test if direct stimulation of uv1 cells is sufficient to induce Ca^2+^ activity, uv1 cells were stimulated using a glass probe, as previously described (Medrano and Collins, 2023). Briefly, a motorized stage was used to move the worm body towards the immobilized probe, varying the distance and time of movement. To create prodding probes, filamented glass capillaries (4”, 1.0 mm outer diameter; World Precision Instruments, 1B100F-4) were pulled using a Sutter P-97 Flaming/Brown type micropipette puller using a custom program with the following parameters: Pressure = 500, Heat = 632 (ramp test value), Pull = 45, Velocity =40, Delay =250.

To perform prodding, a single worm was picked into a small drop of halocarbon oil on a dried 2% agarose pad on a 25 by 60 mm #1 coverslip to immobilize the worm. This immobilized worm then was placed onto an inverted Zeiss Axio Observer.Z1 fluorescence microscope for imaging. A Narishige coarse (model MMN-1) and fine (model MMO-203) micromanipulator, or a Sutter motorized micromanipulator, was used to position the tip of the probe next to the vulva of the worm. The first stimulation was delivered precisely 20 s after the onset of synchronous brightfield and fluorescence recording. A second prodding was performed two minutes after the first stimulation while recording uv1 fluorescence.

### Optogenetics

Optogenetics experiments in animals expressing Channelrhodopsin-2 in the vulval muscles were performed as previously described. To make NGM plates with all-trans retinal (ATR) for optogenetics experiments, ATR was added into pre-warmed OP50 *E. coli* with a final concentration of 0.4 mM (Kopchock et al., 2021). Then 200 µL of culture was seeded onto NGM agar plates. Late L4 worms were staged onto plates with or without ATR 24 h prior to the experiment. Ratiometric Ca^2+^ imaging in freely behaving animals was performed as described previously (Ravi et al., 2018b) with the same 470 nm blue light used to both activate ChR2 and excite GCaMP5 fluorescence. The brightfield and fluorescence image acquisition was synchronized at 20 fps, and each recording lasted 2-3 mins.

### Pharmocological assays

<u>Nemadipine:</u> Nemadipine-B (Toronto Research Chemicals Inc.) was dissolved in DMSO at 10 mM, and 10µL or 30µL was added to ∼4 ml melted NGM solution, generating 25 µM or 75 µM nemadipine plates. Control plates containing an equivalent volume of DMSO were made on the same day (0.25% DMSO final concentration). Plates were seeded with a 10-fold concentrated OP50 *E. coli* bacteria culture one day prior to the experiment. Worms were placed onto nemadipine or DMSO plates for at least 2 h prior to prodding experiments but no longer than 3 h to prevent excess egg accumulation.

<u>Floxuridine:</u> Worms were sterilized with floxuridine (FUDR) as previously described (Collins et al., 2016; Ravi et al., 2018a). Briefly, L4 animals were staged onto NGM plates seeded with OP50. 100 µL of floxuridine (10 mg/mL) was added dropwise onto the staged worms and OP50, and animals were incubated at 20 °C for 24 h. Animals were then visually inspected for lack of embryo accumulation prior to mounting for Ca^2+^ recordings.

Histamine: NGM plates containing 5 mM histamine were prepared as described (Pokala et al., 2014) and seeded with OP50 *E. coli* bacteria. To induce acute silencing, worms were incubated onto histamine-containing plates 2-3 h before the experiment.

### Statistical analyses

Quantitative ratiometric analysis was performed after importing Ca^2+^ recordings into Volocity and GCaMP5/mCherry mean ratio measurements were calculated. Prodding-induced Ca^2+^ transients were analyzed for following parameters: peak amplitude, time to peak, and time to decay. Individual Ca^2+^ transient peak amplitude was analyzed by MATLAB script as described (Collins et al., 2016), and each peak was confirmed by visual inspection of GCaMP5/mCherry ratio traces and the original fluorescence recordings. Time to peak was defined as time interval between the triggered start of prodding and when the uv1 Ca^2+^ transient reached its peak value. Time to decay was defined as the time when the uv1 Ca^2+^ transient peak amplitude fell to 37%. In experiments involving freely moving animals, the peak threshold for calling a uv1 Ca^2+^ transient was 0.2 ΔR/R or greater. Weaker Ca^2+^ transients (<0.2 ΔR/R) were operationally considered as to have not elicited a transient (no transient). For prodding experiments, events <0.4 ΔR/R after direct stimulation were considered as not to have elicited a Ca^2+^ transient.

Statistical analyses were performed using Prism 8 (GraphPad). Ca^2+^ peak amplitude, time to peak, time to decay were compared with one-way ANOVA with Bonferroni correction for multiple comparisons, and error bars indicate 95% confidence intervals for the mean value. p<0.05 interactions were considered significant and are labeled with an asterisk. Percentage of Ca^2+^ responses was compared by Fisher’s exact test, and error bars indicate the 95% confidence intervals for the proportion.

